# Structural basis for MurJ inhibition by phage lysis protein Sgl^PP7^ suggesting convergence

**DOI:** 10.1101/2025.10.15.682725

**Authors:** Kaito Hosoda, Hidetaka Kohga, Sixian Wu, Hiroyuki Tanaka, Hideki Shigematsu, Ryoji Miyazaki, Tomoya Tsukazaki

## Abstract

Some bacteriophages encode lysis proteins that inhibit essential bacterial processes, the elucidation of which is valuable for developing antibacterial strategies against drug-resistant pathogens. We determined the cryo-EM structure of the complex between the essential *E. coli* lipid II flippase MurJ and a phage lysis protein, Sgl^PP7^. MurJ was locked in an outward-facing conformation by Sgl^PP7^, similar to the MurJ/Lys^M^ complex; however, distinct interactions suggest convergent evolution among phage lysis proteins.

## Introduction

Bacteriophages induce host cell lysis by utilizing proteins encoded in their genomes. Single-gene lysis (Sgl) proteins, encoded by single-stranded RNA/DNA phages, induce host cell lysis^1–3^. In gram-negative bacteria, the peptidoglycan layer in the periplasm, located between the inner and outer membranes, is essential for cell integrity. Lys^M^, a 37-residue Sgl of *phage M*, induces cell lysis by targeting MurJ, the essential membrane protein that flips peptidoglycan precursor lipid II from the cytoplasm to the periplasm^4^. Structural studies have shown that MurJ, which consists of 14 transmembrane helices divided into an N-lobe (TM1–6), a C-lobe (TM7–12), and TM13–14, transports lipid II via alternating inward- and outward-facing conformations, resembling MATE transporters^5–7^ (Fig. 1a). Furthermore, cryo-EM structural analysis of the MurJ/Lys^M^ complex (JM complex) revealed that Lys^M^ wedges into the outward-facing conformation of MurJ, preventing the conformational changes as well as lipid II transport, which results in cell lysis^8^.

**Figure 1.**
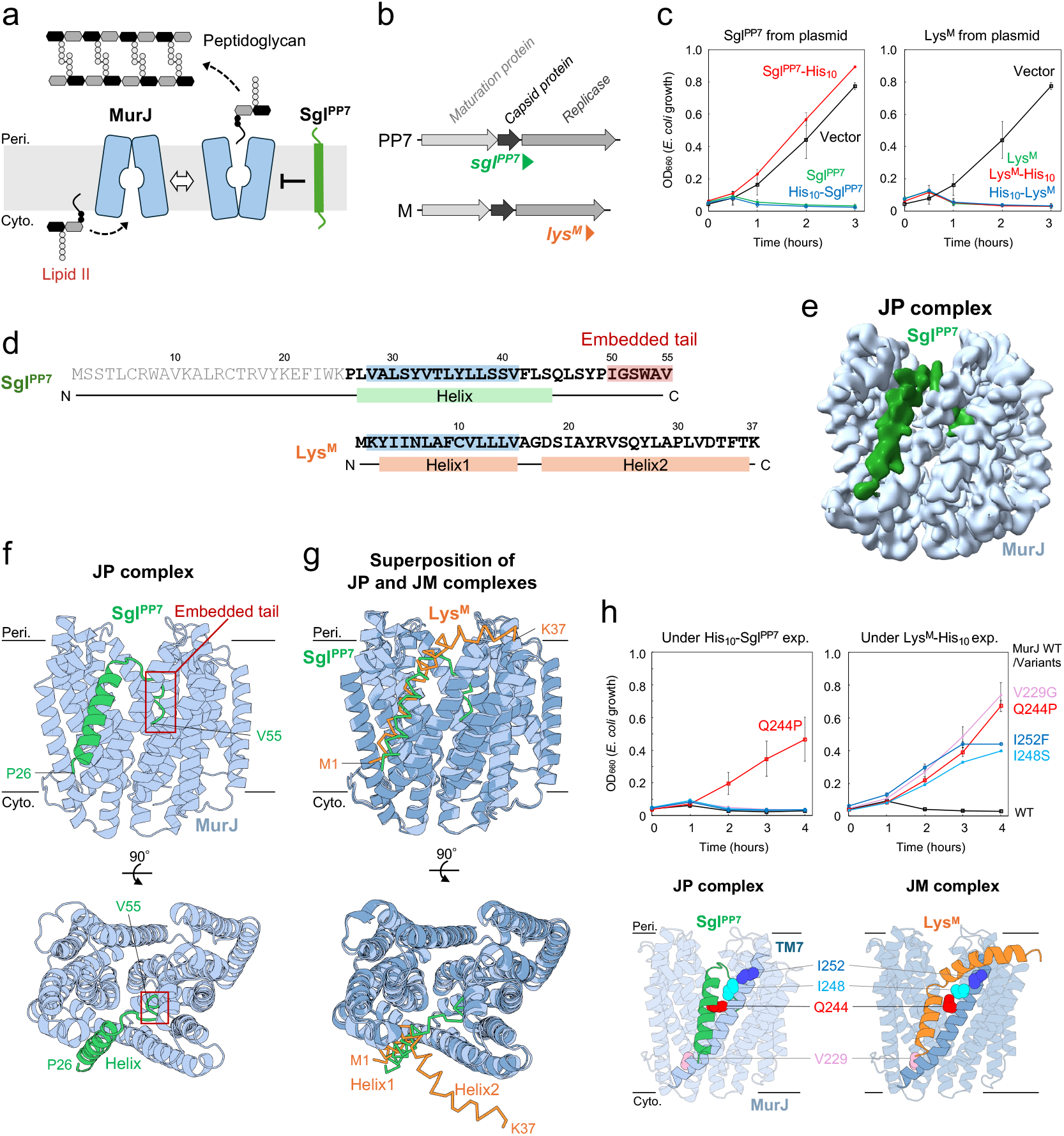
Distinct mode of MurJ inhibition by Sgl^PP7^ compared to Lys^M^. **a**, Inhibition of MurJ-mediated lipid II transport by Sgl^PP7^. **b**, Genomic locations of the single-gene lysis protein genes *sgl*^*PP7*^ (green) in phage PP7 and *lys*^*M*^ (orange) in phage M. **c**, Lytic activities of Sgl^PP7^ and Lys^M^. Growth curves of *E. coli* after overexpression of Sgl^PP7^ and Lys^M^ with or without a His_10_ tag from plasmids induced at 0 min. **d**, Amino acid sequences and secondary structures of Sgl^PP7^ and Lys^M^. Disordered residues of Sgl^PP7^ are shown in grey. Structurally proximate regions between Sgl^PP7^ and Lys^M^ are highlighted in blue. **e, f**, Cryo-EM structure of the JP (MurJ/Sgl^PP7^) complex. (e) Cryo-EM map (contour level, 0.0348) and (f) cartoon models. **g**, Structural superposition of the JP and JM complexes (PDB ID: 9UKV) with MurJ molecules. MurJ and Sgl^PP7^ in the JP complex are coloured as in (f). MurJ and Lys^M^ in the JM complex are shown in deeper blue and orange, respectively. **h**, Activity of Sgl^PP7^ against Lys^M^-resistant MurJ mutants. Growth curves of *E. coli* expressing Lys^M^-resistant MurJ under Sgl^PP7^ or Lys^M^ induction (induction at 0 min, top). The positions of Lys^M^-resistance mutations mapped onto the JP and JM complexes (bottom).

Recently, genetic screening suggested that Sgl^PP7^, a 55-residue Sgl of *Pseudomonas aeruginosa* phage PP7, also targets MurJ^9^. However, compared with Lys^M^, Sgl^PP7^ is encoded at a distinct genomic position and shows no amino acid sequence similarity (Fig. 1b). These findings indicate convergent evolution, wherein evolutionarily unrelated phages have independently acquired the ability to target MurJ. Thus, understanding Sgl mechanisms is important for the development of antibiotics inhibiting MurJ. In this study, we determined the cryo-EM structure of the MurJ/Sgl^PP7^ complex (JP complex), demonstrating that Sgl^PP7^ locks MurJ in the outward-facing conformation through interactions distinct from Lys^M^.

### Results and Discussion

We evaluated the toxicity of Sgl^PP7^ in *E. coli* cells. The induction of untagged or N-terminal His_10_-tagged Sgl^PP7^ caused cell lysis, whereas the C-terminal His_10_-tagged Sgl^PP7^ did not exhibit lytic activity (Fig. 1c, left). In contrast, Lys^M^, which shows little sequence similarity to Sgl^PP7^ (Fig. 1d), retained lytic activity when fused with either the N- or C-terminal tag (Fig. 1c, right). These results indicate that although both Lys^M^ and Sgl^PP7^ are known to inhibit MurJ^9^, Sgl^PP7^ may block MurJ through a mechanism distinct from that of Lys^M^. We therefore purified the MurJ/Sgl^PP7^ (JP) complex using the N-terminal His_10_-tagged Sgl^PP7^ and determined its cryo-EM structure at an overall resolution of 3.04 □(Fig. 1e, Supplementary Fig. 1, Supplementary Table 1).

In the complex structure, MurJ adopts an outward-facing conformation, and Sgl^PP7^ is positioned along the intramembrane cleft of MurJ (Fig. 1e), indicating that Sgl^PP7^ locks MurJ in the outward conformation. The N-terminal residues of Sgl^PP7^ (residues 1–25) are disordered, and the structure consists of a helix that primarily interacts with TM7 of MurJ and a C-terminal tail (Fig. 1e, f; Supplementary Fig. 1e). The tail (residues 50–55) inserts deeply into the positively charged internal cavity on the C-lobe side of MurJ, formed by TM7, TM8, and TM10, and is designated the “embedded tail” (Fig. 1f). This embedded tail likely plays a central role in the interaction with MurJ. This structural feature is consistent with the loss of activity observed for the C-terminal His-tagged Sgl^PP7^. Notably, AlphaFold failed to predict the correct positioning of Sgl^PP7^ on MurJ and showed low confidence in this region, underscoring the importance of the experimentally determined JP complex structure (Supplementary Fig. 2). Moreover, AlphaFold MurJ models adopt an inward-facing conformation, inconsistent with our experimentally determined outward-facing conformation.

In both the JM^8^ and JP complexes, MurJ molecules adopts the outward-facing conformation, with an RMSD of 0.789 Å for the Cα atoms indicating that both MurJ molecules essentially adopt the same outward-facing conformation (Fig. 1g). Lys^M^ consists of Helix1, located near TM7 and TM2, and Helix2, aligned along the membrane surface^8^, whereas the helix of Sgl^PP7^ is located in a similar position to that of Helix1 but takes on a distinct orientation with a different angle relative to TM7. The C-terminal Helix2 of Lys^M^ is located on the exterior of MurJ. In contrast, the embedded tail of Sgl^PP7^ occupies the internal cavity of the C-lobe, creating a markedly different interaction interface with MurJ (Fig. 1g). Importantly, this comparison suggests that although Sgl^PP7^ and Lys^M^ both stabilize MurJ in the outward-facing conformation, they have evolved distinct molecular strategies. Here, we tested the activity of Sgl^PP7^ against four Lys^M^-resistant MurJ mutants^4^. Only cells carrying the Q244P mutation survived upon Sgl^PP7^ induction, although the other Lys^M^-resistant mutants were sensitive to SglPP7 (Fig. 1h). These differential sensitivities support the notion that Lys^M^ and Sgl^PP7^ inhibit MurJ through distinct binding interfaces.

We addressed the importance of the embedded tail of Sgl^PP7^ by mutational analysis. Successive C-terminal truncations revealed that residues up to residue 53 are essential for Sgl^PP7^ function (Fig. 2c, left; Supplementary Fig. 3a). Next, we examined the lytic activity of several mutations at W53, which inserts into the C-lobe (Fig. 2a, b; 2c, middle; Supplementary Fig. 3b). The mutant W53A retained full activity, W53G, W53C, and W53S showed slightly reduced activity, and W53K and W53R lost lytic activity. These results suggest that W53 is important for stabilizing the interaction with MurJ, although this effect may be partially compensated by neighbouring residues, including residues 54–55. We then examined the same W53 variants in the truncated Sgl^PP7^(1–53) mutant, which retains minimal activity, and all the variants showed weakened lytic activity (Fig. 2c, right; Supplementary Fig. 3c). These results suggest that the C-terminal region is an important determinant of Sgl^PP7^ function, with W53 serving as a key residue within this region.

**Figure 2.**
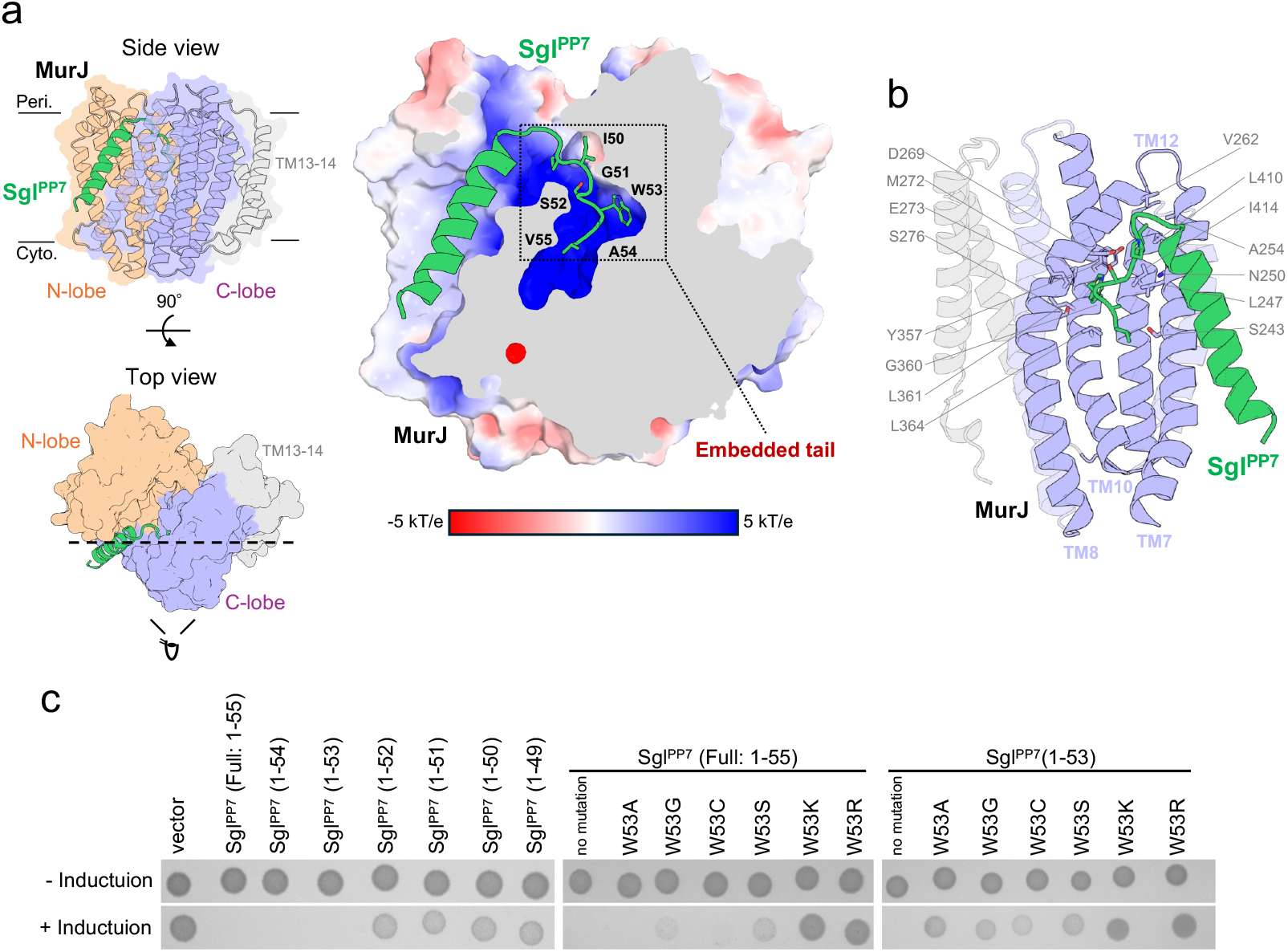
Embedded tail of Sgl^PP7^ fitted into the C-lobe cavity of outward-facing MurJ. **a**, Cutaway model of the JP complex along the dashed line. (Left) The N-lobe, C-lobe, and TM13–14 of MurJ are shown in orange, purple, and grey, respectively. (Right) Cutaway of MurJ is shown as an electrostatic potential surface, ranging from -5 kT/e (red) to +5 kT/e (blue). Residues 26–48 of Sgl^PP7^ are shown in cartoon representation and residues 49–55 are shown in stick representation. **b**, Embedded tail surrounded by TMs 7, 8, and 10 of MurJ. The C-lobe is viewed from the N-lobe side, with the N-lobe omitted for clarity. MurJ residues positioned within 4 Å of Sgl^PP7^ are shown as sticks. **c**, Lytic activity of C-terminally truncated mutants (left) and W53 substitution mutants in full-length or truncated Sgl^PP7^(1–53) (middle or right, respectively). *E. coli* cells harbouring vector plasmids or plasmids encoding Sgl^PP7^ mutants were grown on agar plates with or without induction.

Our findings demonstrate that the unique embedded tail of Sgl^PP7^ inserts into the internal cavity of MurJ, locking it in the outward-facing conformation (Fig. 1g, h). In this way, Sgl^PP7^ physically disrupts the lipid II flippase cycle of MurJ, halting the peptidoglycan biosynthesis. Although both Sgl^PP7^ and Lys^M^ stabilize MurJ in the same outward-facing conformation, their helix orientations and binding interfaces are inherently different. Our structural studies show that evolutionarily unrelated phages have convergently acquired the ability to inhibit MurJ. Beyond these two examples, unidentified phage proteins could also target MurJ through distinct mechanisms, highlighting the importance of further exploring the diversity of phage-encoded lysis strategies, which may ultimately inform the development of next-generation antibacterial agents.

## Methods

### Bacterial Strains and Plasmids

The *E. coli* strains and plasmids used in this study are listed in Supplementary Table 2. Details of the plasmids used are described in the Plasmid Construction section.

### Plasmid Construction

A DNA fragment containing the *sgl*^*PP7*^ gene, which was codon optimized for *E. coli*, was synthesized de novo (Integrated DNA Technologies, IDT). The *sgl*^*PP7*^ fragment was cloned into XbaI-digested pBAD33^10^ using In-Fusion HD Cloning Kit (Takara Bio). The resulting plasmid was designated pKG382. For pKH32 (pBAD18^10^ encoding *his*_*10*_*-sgl*^*PP7*^), the *his*_*10*_*-sgl*^*PP7*^ fragment was PCR-amplified from pKG382. The PCR product was cloned into inverse PCR-linearized pBAD18 using In-Fusion HD Cloning Kit (Takara Bio). Mutant plasmids were constructed using a site-directed mutagenesis.

### Media and Bacterial Cultures

*E. coli* cells were grown at 37°C in LB medium (Nacalai Tesque). ampicillin (Amp) (100 µg/mL), chloramphenicol (Cm) (20 µg/mL), and spectinomycin (Spc) (50 µg/mL) were added as appropriate for plasmid-bearing cell growth and transformant selection. Bacterial growth was monitored with a Mini photo 518R (660 nm; TAITEC Co., Japan).

### Lysis assay

*E. coli* BL21 Δ*recA* (DE3) cells carrying a plasmid encoding one of the Sgl^PP7^ derivatives were grown overnight in LB medium supplemented with the appropriate antibiotics. The cultures were diluted 1:10,000 in phosphate-buffered saline and spotted onto LB agar plates supplemented with 0.2% (w/v) L-arabinose and the necessary antibiotics. The plates were incubated at 37°C for 22 h and subsequently imaged.

### Sample preparation of the MurJ/Sgl^PP7^ complex

*E. coli* BL21 Δ*recA* (DE3) cells harbouring the plasmids pKG537, pKH32, and pRM1210, encoding MA-*Ec*MurJ_2-511_ (from *E. coli* JCM20135)-3×FLAG, His_10_-Sgl^PP7^, and *Bs*Amj, respectively, were cultured in LB medium (Lennox, Nacalai) supplemented with 100 µg/mL Amp, 20 µg/mL Cm, and 50 µg/mL Spc at 37°C. Protein expression was induced with 1 mM isopropyl β-D-thiogalactopyranoside (IPTG) and 0.2% (w/v) L-arabinose. Protein purification was performed as described previously^8^. The membrane fractions were isolated by ultracentrifugation and solubilized in buffer containing 2% n-dodecyl β-maltoside (DDM), 0.02% lauryl maltose neopentyl glycol (LMNG), and 0.002% cholesterol hydrogen succinate (CHS). The protein was subsequently purified using Ni-NTA agarose resin (QIAGEN), followed by a Superose 6 Increase 10/300 GL column (Cytiva). The MurJ/Sgl^PP7^ complex was finally purified in buffer (20 mM Tris-HCl (pH 8.0), 300 mM NaCl, 0.005% LMNG, 0.0005% CHS, and 0.1 mM PMSF) and concentrated to ∼20 mg/mL using an Amicon Ultra 30 K NMWL (MERCK). Nanodisc reconstitution was performed as described previously^8^. The purified MurJ/Sgl^PP7^ complex was mixed with MSP1D1^11^ and *E. coli* phospholipids (Avanti) at a molar ratio of 1.4:1:120 in buffer (20 mM Tris-HCl (pH 8.0), 300 mM NaCl, 0.005% LMNG, 0.0005% CHS, and 0.1 mM PMSF). The mixture was incubated at 4°C for 1 h. Then, the detergent was removed by Bio-Beads SM2 (Bio-Rad). The resulting nanodiscs were further purified using a Superdex 200 Increase 10/300 GL column. preequilibrated with buffer (20 mM Tris-HCl (pH 8.0), 300 mM NaCl, and 0.1 mM PMSF). The MurJ/Sgl^PP7^-reconstituted nanodisc was concentrated using an Amicon Ultra 50 K NMWL.

### Cryo-EM data collection and processing

Quantifoil holey carbon grids (Cu R1.2/R1.3, 300 mesh) were glow-discharged at 7 Pa and 10 mA for 10 s using a JEC-3000FC sputter coater (JEOL) before sample application. A 3 μL aliquot of 2.8 mg/mL MurJ/Sgl^PP7^ complex-reconstituted nanodiscs was applied onto the grid, blotted for 2 s at 100% humidity, 8°C, and blot force 10, and plunged into liquid ethane using a Vitrobot Mark IV (Thermo Fisher Scientific).

The cryo-EM dataset was collected on a CRYO ARM 300 transmission electron microscope (JEOL) operated at an accelerating 300 kV, equipped with a cold-field emission gun, an in-column Omega-type energy filter, and a Gatan K3 direct electron detector (Gatan) at SPring-8. Images were collected at a nominal magnification of 60,000×, corresponding to a calibrated pixel size of 0.752 Å/pixel, 50 frames per image with a total exposure dose of 50 e^−^/Å^2^ using a SerialEM^12^.

Data processing was performed using CryoSPARC v4.6.2 software^13^. The cryo-EM data-processing workflow for the MurJ/Sgl^PP7^ complex is shown in Supplementary Figure 1a. A total of 17,650 movies were aligned using patch motion correction, and the contrast transfer function (CTF) parameters were estimated using the Patch CTF estimation. The particles were automatically picked using the Blob picker in CryoSPARC among 200 of the 17,650 micrographs. Particles were extracted with downsampling via Fourier cropping to a pixel size of 1.504 Å and subjected to 2D classification, generating 2D class averages. Using these 2D class averages as templates, particles were repicked using the Template Picker from 17,650 micrographs, yielding 7,554,685 particles. The repicked particles were extracted with downsampling via Fourier cropping to the pixel size of 1.504 Å and subjected to five rounds of 2D classification (K = 350, 350, 300, 250, 250). After 2D classification, approximately 1,774,539 high-quality particles were selected and re-extracted to 0.752 Å/pixel. These selected particles were subjected to ab initio 3D reconstruction (K = 4), generating initial 3D models. After two rounds of 2D classification, the 3,749,310 particles were then subjected to five rounds of heterogeneous refinement (K = 4, 4, 4, 4, 4) using the initial models as references, removing junk particles. The best 3D class, containing 386,409 particles, was re-extracted to 0.752 Å/pixel and refined using Non-Uniform (NU) refinement. These particles were subjected to reference-based motion correction and 2D classification (K = 100). After 2D classification, 347,457 cleaned and polished particles were subjected to NU refinement. These particles were subsequently subjected to global CTF refinement, local CTF refinement, NU refinement, and model building. The resulting particles were used for local refinement using a mask created around the region of interest with a radius of 12 Å. These processes yielded a 3D cryo-EM map with an estimated overall resolution of 3.04□Å. Local resolution was estimated using the Local Resolution Estimation job in CryoSPARC.

### Model building and refinement

The Sgl^PP7^ structure was predicted using AlphaFold3^14^. The predicted Sgl^PP7^ model and an EcMurJ outward model from the MurJ/Lys^M^ complex structure (PDB ID: 9UKV) were docked into the cryo-EM map using UCSF ChimeraX. The model was manually modified in Coot^15^, and further refinement was conducted using phenix.real_space_refine in PHENIX^16^. Data processing and refinement statistics are provided in Supplementary Figure 1 and Supplementary Table 1. The final molecular model and cryo-EM map were visualized using PyMOL (https://pymol.org/) or UCSF ChimeraX^17^.

## Acknowledgements

We thank Kayo Abe for secretarial assistance, Kunihito Yoshikaie for experimental advice, and the scientists of SPring-8 Structural Biology Beamlines for helping with data collection. The cryo-EM experiments were performed at SPring-8 with the approval of the Japan Synchrotron Radiation Research Institute (Proposal Nos. 2024A2742, 2024A2759, 2025A2725, and 2025A2765). We gratefully acknowledge Nara Institute of Science and Technology for allowing access to its facilities on weekends and public holidays, which enabled part of this work as voluntary, unpaid “self-training” (Jikokensan) research. This work was supported by the JSPS/MEXT KAKENHI (Grant Nos. JP25K18422 and JP23K14146 to H.K.; JP25K00267, JP24KK0138, JP22K15061, and JP22H05567 to R.M.; and JP25K02226, JP22H02567, JP22H02586, JP21H05155, JP21H05153, and JP21KK0125 to T.T.); private research foundations (The Chemo-Sero-Therapeutic Research Institute, ONO Medical Research Foundation and Takeda Science Foundation to T.T.; and the Institute for Fermentation (Osaka) to H.K., R.M. and T.T.); and JST SPRING (Grant No. JPMJSP2140 to K.H.). This research was partially supported by Platform Project for Supporting Drug Discovery and Life Science Research (Basis for Supporting Innovative Drug Discovery and Life Science Research (BINDS)) from AMED under grant number JP24ama121001.

## Author contributions

H.K. and T.T. conceptualized the study. K.H., H.K., S.W., H.T., and R.M. performed the biological analyses. K.H., H.K., H.S., and T.T. determined the structure. K.H., H.K., and T.T. wrote the manuscript. H.K., T.M., and T.T. supervised the study.

## Data and Materials Availability

The coordinates and the electron potential maps for the JP complex structure have been deposited in PDB and EMDB under accession codes: PDB: 9X2N and EMBL-EBI: 66480, respectively.

## Supplementary materials

Figs. S1–S3

Tables S1–2

**Supplementary Figure 1. Cryo-EM data processing workflow for the JP complex**.

**Supplementary Figure 2. AlphaFold-predicted models of MurJ/Sgl**^**PP7**^ **complex**.

**Supplementary Figure 3. Uncropped spot assay images corresponding to Figure 2c**.

**Supplementary Table 1. Cryo-EM data collection and refinement statistics**.

**Supplementary Table 2. Plasmids and strains used in this study**.

