## Supplemental figs and tables for "Structural basis for MurJ inhibition by phage lysis protein Sgl^PP7^ suggesting convergence"

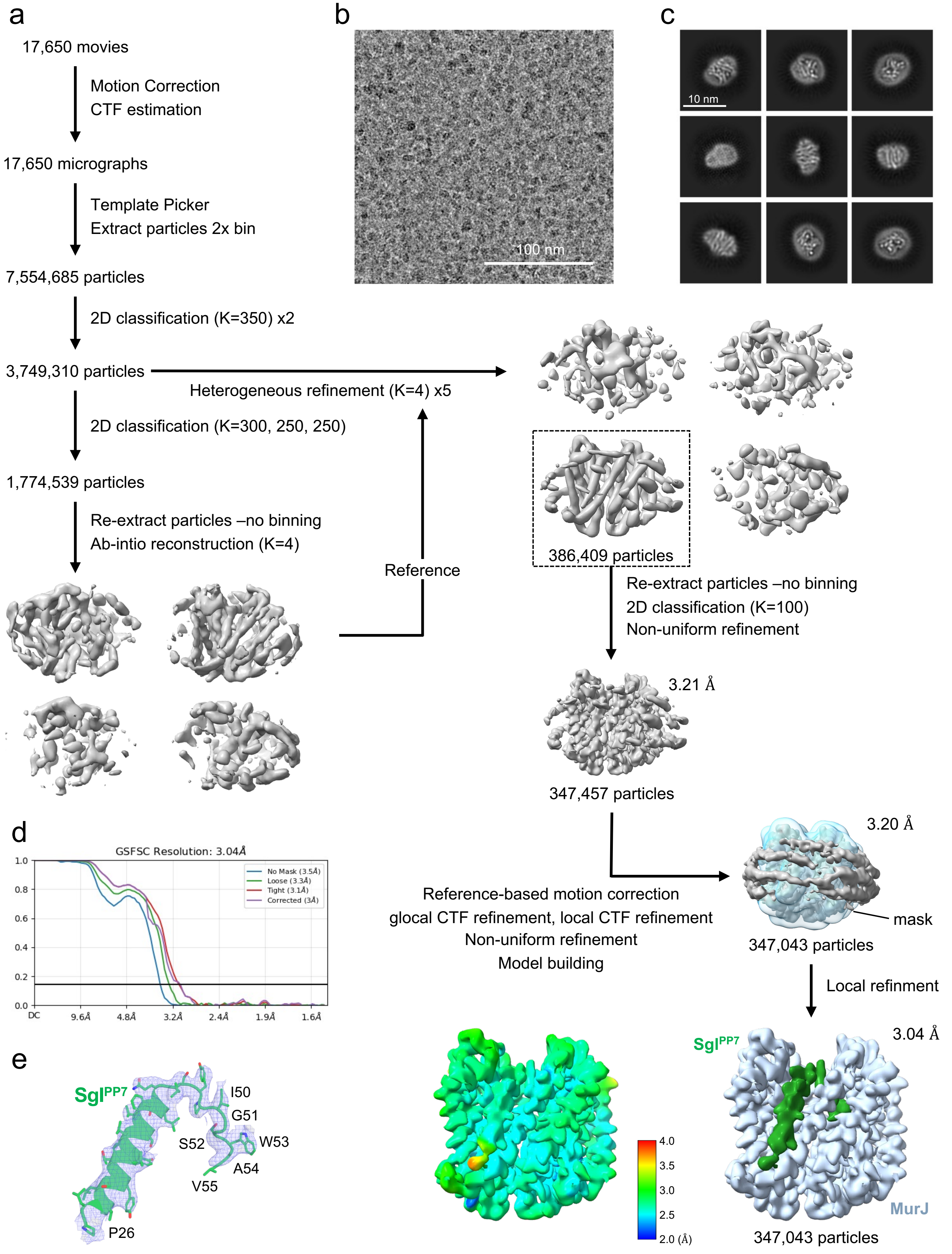


**Supplementary Figure 1. Cryo-EM data processing workflow for the JP complex.**
**a,** Image processing workflow for the JP complex. The local resolution of the reconstructed map was estimated using CryoSPARC. **b,** Representative electron micrograph of the nanodisc-reconstituted JP complex. **c,** 2D class averages of the JP complex. **d,** Gold-standard FSC curve used for global resolution estimation within CryoSPARC. **e,** Cryo-EM map (displayed at 6σ) of $Sgl^{PP7}$ and its model.

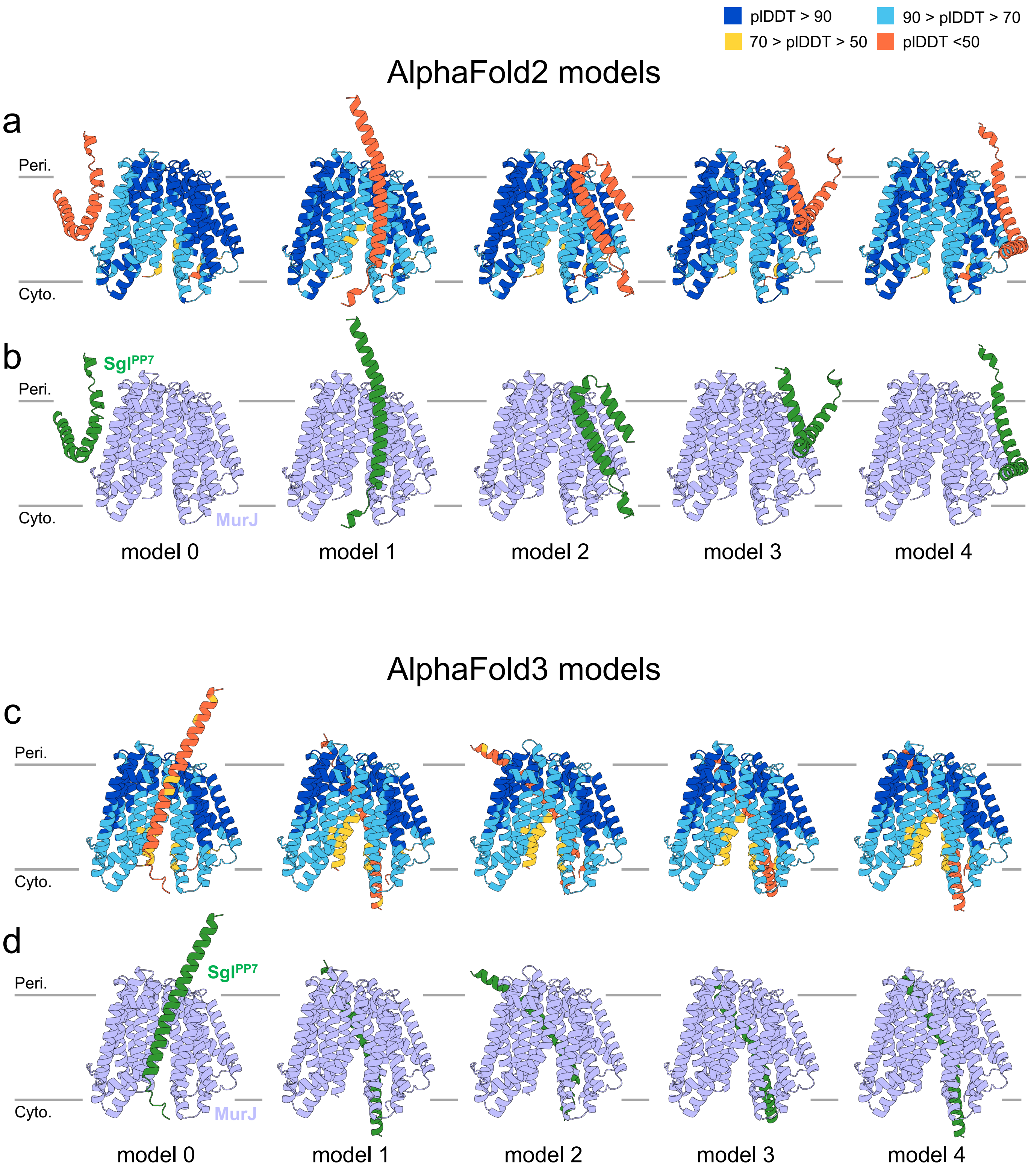


**Supplementary Figure 2. AlphaFold-predicted models of MurJ/Sgl$^{PP7}$ complex.**
MurJ/Sgl$^{PP7}$ models were generated using AlphaFold2 (colabfold 1.5.2) and AlphaFold3 (https://alphafoldserver.com/). **a** and **c,** Predicted models colour-coded by per-residue pLDDT scores. b and **d,** Predicted models coloured light blue (MurJ) and green (Sgl$^{PP7}$).

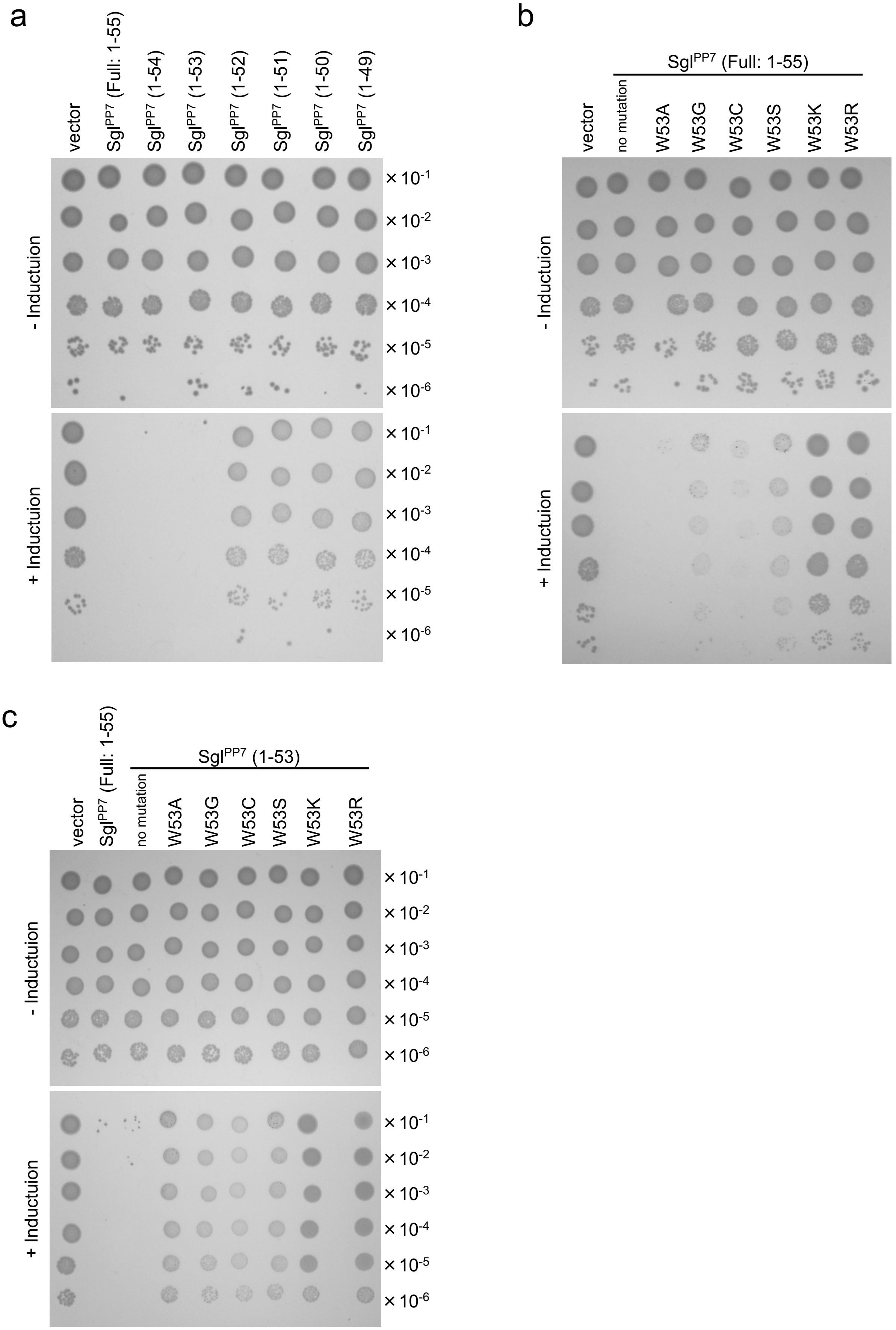


**Supplementary Figure 3. Uncropped spot assay images corresponding to Figure 2c.**
**a,** Original images in Fig. 2c (left) showing the cell lytic activities of the C-terminal-truncated His$_{10}$-Sgl$^{PP7}$ mutants. **b,** Original images in Fig. 2c (middle) showing the cell lytic activities of the His$_{10}$-Sgl$^{PP7}$ W53 substitution mutants. **c,** Original images in Fig. 2c (right) showing the cell lytic activities of the His$_{10}$-Sgl$^{PP7}$ (1-53) W53 substitution mutants.

**Table S1. Data collection and refinement statistics.**

| | *Ec*MurJ/Sgl$^{PP7}$-ND (EMD-66480, PDB ID: 9X2N) |
|---|---|
| **Data collection and processing** | |
| Magnification | x60,000 |
| Voltage (kV) | 300 |
| Electron exposure (e–/Å$^2$) | 50 |
| Defocus range (μm) | -1.4 to -1.6 |
| Pixel size (Å) | 0.752 |
| Symmetry imposed | C1 |
| Initial particle images (no.) | 7,554,685 |
| Final particle images (no.) | 347,457 |
| Map resolution (Å) | 3.04 |
| FSC threshold | 0.143 |
| Map resolution range (Å) | 32.26 to 2.00 |
| **Refinement** | |
| Initial model used (PDB code) | 9UKV and AlphaFold model |
| Model composition | |
| Non-hydrogen atoms | 4121 |
| Protein residues | 540 |
| Ligands | 0 |
| *B* factors (Å$^2$) | |
| Protein | 206.17 |
| Ligand | 0 |
| R.m.s. deviations | |
| Bond lengths (Å) | 0.005 |
| Bond angles (°) | 0.830 |
| **Validation** | |
| MolProbity score | 1.74 |
| Clashscore | 8.25 |
| Poor rotamer (%) | 2.07 |
| Ramachandran plot | |
| Favored (%) | 97.77 |
| Allowed (%) | 2.23 |
| Disallowed (%) | 0 |

**Table S2. Plasmids and Strains used in this study.**

| Plasmids | Vector | Encoded gene and description | Reference or source |
| --- | --- | --- | --- |
| pBAD33 | - | Expression vector, $P_{araBAD}$, $Cm^R$ | *(10)* |
| pKK568 | pBAD33 | $lys^M$ | *(8)* |
| pKG265 | pBAD33 | $lys^M$-$his_{10}$ | This study |
| pKG360 | pBAD33 | $his_{10}$-$lys^M$ | This study |
| pKG382 | pBAD33 | $sgl^{PP7}$ | This study |
| pKG398 | pBAD33 | $sgl^{PP7}$-$his_{10}$ | This study |
| pKG404 | pBAD33 | $his_{10}$-$sgl^{PP7}$ | This study |
| pBAD18 | - | Expression vector, $P_{araBAD}$, $Amp^R$ | *(10)* |
| pKH32 | pBAD18 | $his_{10}$-$sgl^{PP7}$ | This study |
| pKH57 | pBAD18 | $his_{10}$-$sgl^{PP7}$(1-54) | This study |
| pKH40 | pBAD18 | $his_{10}$-$sgl^{PP7}$(1-53) | This study |
| pKH42 | pBAD18 | $his_{10}$-$sgl^{PP7}$(1-52) | This study |
| pKH59 | pBAD18 | $his_{10}$-$sgl^{PP7}$(1-51) | This study |
| pKH45 | pBAD18 | $his_{10}$-$sgl^{PP7}$(1-50) | This study |
| pKH48 | pBAD18 | $his_{10}$-$sgl^{PP7}$(1-49) | This study |
| pKG743 | pBAD18 | $his_{10}$-$sgl^{PP7}$(W53A) | This study |
| pKH69 | pBAD18 | $his_{10}$-$sgl^{PP7}$(W53G) | This study |
| pKH72 | pBAD18 | $his_{10}$-$sgl^{PP7}$(W53C) | This study |
| pKH75 | pBAD18 | $his_{10}$-$sgl^{PP7}$(W53S) | This study |
| pKH70 | pBAD18 | $his_{10}$-$sgl^{PP7}$(W53K) | This study |
| pKH68 | pBAD18 | $his_{10}$-$sgl^{PP7}$(W53R) | This study |
| pKH81 | pBAD18 | $his_{10}$-$sgl^{PP7}$(1-53, W53A) | This study |
| pKH77 | pBAD18 | $his_{10}$-$sgl^{PP7}$(1-53, W53G) | This study |
| pKH78 | pBAD18 | $his_{10}$-$sgl^{PP7}$(1-53, W53C) | This study |
| pKH79 | pBAD18 | $his_{10}$-$sgl^{PP7}$(1-53, W53S) | This study |
| pKH80 | pBAD18 | $his_{10}$-$sgl^{PP7}$(1-53, W53K) | This study |
| pKH76 | pBAD18 | $his_{10}$-$sgl^{PP7}$(1-53, W53R) | This study |
| pSTV28 | - | Expression vector, $P_{lac}$, $Cm^R$ | Takara Bio |
| pKG537 | pSTV28 | *EcmurJ-3xflag* | *(8)* |
| pKG50 | pTSP1 | *EcmurJ* | This study |
| pKG59 | pTSP1 | *EcmurJ* (V229G) | This study |
| pKG678 | pTSP1 | *EcmurJ* (Q244P) | This study |
| pKG681 | pTSP1 | *EcmurJ* (I248S) | This study |
| pKG646 | pTSP1 | *EcmurJ* (I252F) | This study |
| pRM1210 | pHM1550 | *Bacillus subtilis amj* | *(8)* |

| Strains | Genotype | Reference or source |
| --- | --- | --- |
| JM109 | *recA1*, *endA1*, *gyrA96*, *thi-1*, *hsdR17*(*rK- mK*+), *e14-* (*mcrA*-), *supE44*, *relA1*, Δ(*lac-proAB*)/F'〔*traD36*, *proAB*+, *lac Iq*, *lacZ*ΔM15〕 | Takara Bio |
| BL21 Δ*recA* (DE3) | *F-, ompT, hsdSB*（*rB-mB-*）*, gal, dcm,* Δ*recA,* Δ*endA* | Nippon gene |
